# Microbial piggy-back: how *Streptomyces* spores are transported by motile soil bacteria

**DOI:** 10.1101/2020.06.18.158626

**Authors:** Alise R. Muok, Dennis Claessen, Ariane Briegel

**Affiliations:** Institute for Biology, Leiden University, Sylviusweg 72, 2333 BE Leiden, The Netherlands Centre for Microbial Cell Biology, Leiden University, Leiden, The Netherlands

## Abstract

Streptomycetes are sessile, soil-dwelling bacteria that produce diverse metabolites that impact plant health and the behavior of microbial communities. Emerging studies have demonstrated that *Streptomyces* spores are distributed through a variety of mechanisms, but it remains unclear how spores are transported to their preferred micro-environments, such as plant roots. Here, we show that *Streptomyces* spores are capable of utilizing the motility machinery of other soil bacteria and are transported on the centimeter scale. Motility assays and microscopy studies reveal that *Streptomyces* spores are transported to plant tissues by interacting directly with the flagella of both gram-positive and gram-negative bacteria. Genetics experiments demonstrate that this form of motility, called piggy-backing, is facilitated by conserved structural proteins present on the surface of *Streptomyces* spores. These results demonstrate that non-motile bacteria are capable of utilizing the motility machinery of other microbes to complete necessary stages of their lifecycle, and that this mode of transport may be ubiquitous in nature.

## Introduction

Bacteria belonging to the genus *Streptomyces* are an integral component of diverse ecosystems and are well-known to produce chemically diverse metabolites, including the vast majority of all clinically-relevant antibiotics^1,2^. Soil-dwelling Streptomycetes, such as *Streptomyces coelicolor*, colonize plant roots and provide the associated plant protection from potential phytopathogens through antibiotic secretion^1^. The symbiosis of Streptomycetes with their plant hosts has been shown to improve plant health and productivity, and thereby provides a potential sustainable solution to increase crop yields^2–5^. The lifecycle of Streptomycetes is complex and involves stages of aerial hyphae formation on the soil surface to produce spores, and spore germination on plant roots to produce filamentous colonies^1^. Immotile *Streptomyces* bacteria distribute their spores over long distances through attachment to insects and nematodes, but it is unclear how they relocate over short distances to their preferred microenvironments such as plant root systems^6,7^. While *Streptomyces* are non-motile, many other soil microbes are motile and can sense plant root exudates via bacterial chemotaxis^8,9^.

Bacterial chemotaxis is the system employed by most motile bacteria and archaea to navigate their environment. It relies on two large cellular machineries: the chemoreceptor array and the flagellum. The chemoreceptor array is a membrane-associated supramolecular protein assembly that is relatively conserved across motile bacteria and archaea^10^. By sensing specific ligands through specialized transmembrane receptors in the arrays, the microbes are able to move toward beneficial compounds (attractants) and away from deleterious ones (repellents) by regulating the direction of rotation of the flagella^11,12^. In nature, this behavior allows pathogenic microbes to efficiently navigate host tissues and allows free-living bacteria to seek their preferred microenvironmental niches^13–15^.

Recent reports have revealed that microbe transport by inter-species interactions can occur between motile and immotile microbes. These studies demonstrate that inter-microbial transport occurs among organisms natively found on abiotic surfaces^16,17^, plant surfaces^18^, and within the soil^19,20^. In some instances, the transportation of human pathogens are facilitated on abiotic surfaces, including non-motile Staphylococcal species that directly adhere to their mobile partners^16^, *Aspergillus fumigatus* spores that interact with the flagella of motile bacteria^20^, and *Legionella pneumophila* that is transported internally by their amoebae hosts^21^.

Here, we demonstrate that spores of the sessile Streptomycetes, such as *Streptomyces coelicolor (Sc)*, are transported by *Bacillus subtilis (Bs)* to their preferred microenvironment. *Sc* and *Bs* are both soil-dwelling bacteria that utilize plant root exudate as a nutritive source^1,14^. Unlike *Sc, Bs* is known to sense and steer the cell’s motility toward plant root exudate^14^. Using microscopy methods, motility assays, and genetics approaches, we demonstrate that *Bs* transports *Sc* spores via direct attachment to *Bs* flagella, a mode of transportation we call ‘piggy-backing’. Piggy-backing is dependent on the conserved rodlin proteins, which form a fibrous outer layer on the spore coat of almost all Streptomycetes, but with a hitherto unclear function^22,23^. These results exemplify that non-motile bacteria are capable of utilizing the motility machinery of other microbes to complete necessary stages of their lifecycle, and that this mode of transport may be ubiquitous in nature.

## Results

### Bacillus subtilis disperses Streptomyces coelicolor spores

Transportation of *Sc* spores by *Bs* was demonstrated by mixing isolated *Sc* spores with a liquid culture of *Bs* followed by inoculation onto an agar swarm plate and incubation. After 5 days, *Sc* colonies are visible on the plate and are only dispersed in the presence of *Bs* in all samples tested (*n* = 10) (Fig. 1A). To demonstrate that the *Sc* spores are being moved by *Bs* cells and not merely ‘floating’ in the expanding *Bs* colony, we conducted identical assays but inoculated the *Sc* spores and *Bs* culture separately and onto different areas of the plate. The resulting *Sc* colonies form streaks across the plate that emanate from the *Bs* inoculation site in a predictable manner in all samples tested (*n* = 10) (Fig. 1B). This experiment was repeated on 12 cm plates and demonstrate that *Sc* spores are dispersed to the edge of the plate, which is 10 cm from the spore innoculation point (*n* = 3) (Fig. S1).

**Fig. 1.**
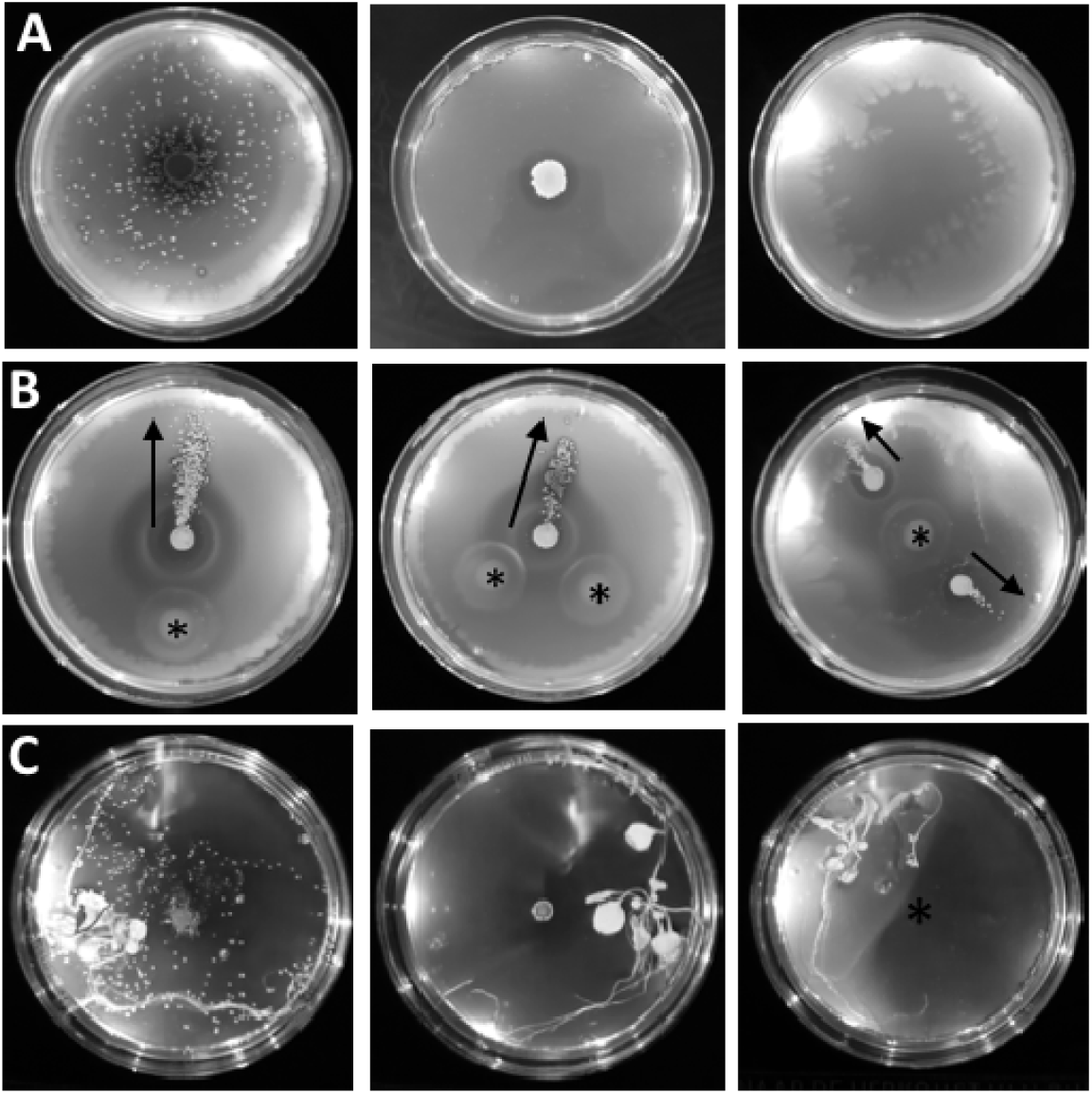
*S. coelicolor* spores are transported by *B. subtilis*. (A) When *Sc* and *Bs* are innoculated on the center of a swarm plate, visible *Sc* colonies (white dots) are apparent and are only dispersed in the presence of motile *Bs*. Left: *Sc* with *Bs*. Middle: *Sc* alone. Right: *Bs* alone. (B) When *Sc* and *Bs* are innoculated in different positions on swarm plates, the *Sc* colonies are dispersed in the swarming direction of the *Bs* cells (black arrows). Asterisks denote the *Bs* innoculation sites. (C) *Bs* moves spores toward plant tissues. Left: *Sc* with *Bs*. Middle: *Sc* alone. Right: *Bs* alone, asterisk denotes the *Bs* innoculation site.

### B. subtilis transports S. coelicolor spores to plant tissues

In nature, *Sc* and *Bs* thrive near plant roots that excrete exudates but only *Bs* can sense and move toward the exudates. We conducted assays to determine if *Bs* can transport *Sc* spores to plant tissues. Assays with the *Bs* strain alone demonstrate that the plates become ‘cloudy’ with *Bs* cells in areas around plant tissues (Fig. 1C). Like previous experiments, the *Sc* spores alone do not exhibit movement unless they are co-inoculated with *Bs* cells, and the dispersed spores preferentially establish colonies near plant tissues in all instances (*n* = 5) (Fig. 1C).

### Spore dispersal occurs through bacterial swarming

*Bs* has two modes of flagellar-mediated motility, swimming and swarming, that occur in liquid environments and on solid surfaces, respectively. When *Bs* senses that it is on a solid surface, it will differentiate into a swarmer cell that has a significant increase in the number of flagella and produces hydrophobic surfactants, such as surfactin, to efficiently spread across the surface^24–26^. Both swimming and swarming cell types are chemotactic^27,28^. In our experiments, we utilized an undomesticated strain of *Bs* (NCIB3610) that can swarm, unlike common laboratory strains that lack the ability to differentiate into swarmer cells and fail to produce surfactin to undergo swarming motility^24,26,29^. To determine if the *Sc* spores are transported by both swarming and swimming motilities, we repeated the experiments with a laboratory cultivated *Bs* strain that is incapable of swarming, and spore transport does not occur on the agar plates^24^ (Fig. S2). Therefore, we conclude that spore transport is accomplished via swarming motility.

### S. coelicolor spores attach to B. subtilis flagella

We utilized several microscopy methods to elucidate a mechanism for *Sc* spore dispersal by *Bs*. Fluorescently-labeled *Sc* spores were imaged under a fluorescence microscope and were immotile as expected (<u>Supplementary Movie 1</u>). However, when *Bs* cells were added to the fluorescent spores, the spores localize near the *Bs* cell poles and are motile (<u>Supplementary Movie 2</u>, Fig. 2A). In some instances, the spores are stationary on the surface of the glass slide and an associated *Bs* cell is seen rotating around the spore (<u>Movie 1</u>). The observed *Bs* cell rotation in these assays is reminiscent of rotations seen in *Bs* cells that have their flagella chemically tethered to a solid surface, whereby the torque generated by the immobilized flagella induces rotation of the cell body^30^. This observation suggests that the *Sc* spores adhere directly to the *Bs* flagella, and therefore effectively mimic a flagellar tether in these instances. To verify that the *Sc* spores do not directly interact with the *Bs* cell body, we imaged a mixture of *Sc* spores and *Bs* cells with a cryo-electron microscope. Like the fluorescence microscopy images, the *Sc* spores are localized near the *Bs* cell poles but do not make direct contact with the *Bs* cell body (Fig. 2B). To confirm that spores directly adhere to the flagella, we utilized a *Bs* ‘mini-cell’ strain (minD::TnYLB) that lacks the excreted material inhibiting their direct visualization. Indeed, when mixed with *Sc* spores, the flagella can be seen co-localizing with the spores in two-dimensional cryo-EM images (Fig. 2C).

**Fig. 2.**
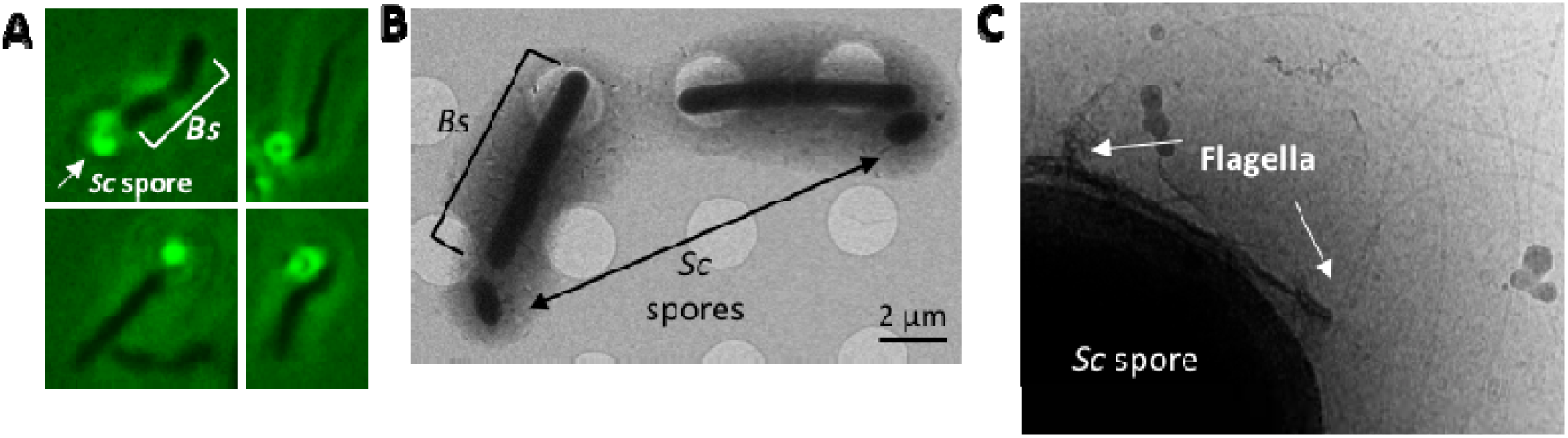
Microscopy methods indicate that *Sc* spores directly adhere to *Bs* flagella. (A) Fluorescence microscopy of dye-labeled spores with unlabeled *Bs* cells demonstrate that the spores localize to the cell poles of *Bs*. (B) Cryo-electron microscopy samples of mixed *Sc* spores and *Bs* cells reveal that the spores do not directly adhere to the *Bs* cell body. (C) Cryo-EM shows the *Bs* flagella colocalize with the *Sc* spore coat.

### Piggy-backing is conserved in Streptomycetes

To determine if spore dispersal by *Bs* also occurs in other *Streptomyces* species, we conducted *Bs* swarm plate assays with *Streptomyces tendae (St), Streptomyces griseus (Sg), Streptomyces scabies (Ss*), and *Streptomyces avermitilis (Sa)*. To quantify spore dispersal, we prepared swarm-plate assays where *Bs* cells were inoculated at the center of the plate (9 cm in diameter) and the isolated spores are inoculated in four equidistant positions around the *Bs* inoculation site. The maximum dispersal distance, which is the distance from the center of the spore inoculation site to the most distant dispersed colony, was measured for each of the four spore samples. The wild-type (WT) *Sc* spores are dispersed by *Bs* in 100% of samples and are moved an average maximum distance of 2.67 +/- 0.43 cm from the initial inoculation point (*n* = 20). Likewise, these assays demonstrate that *St* (*n* = 12), *Sg* (*n* = 8), and *Ss* (*n* = 8) spores are dispersed in 100 % of samples to similar distances as WT *Sc* spores. However, the *Sa* spores are dispersed at significantly shorter distances (*n* = 12) (Fig. 3A,B) and are dispersed in 83 % of the samples. As the last common ancestor of *Sc* and *Sg* existed more than 200 million years ago and both species are capable of piggy-backing, these data suggest that the ancestor also possessed this dispersal mechanism and it remained conserved.

**Fig. 3.**
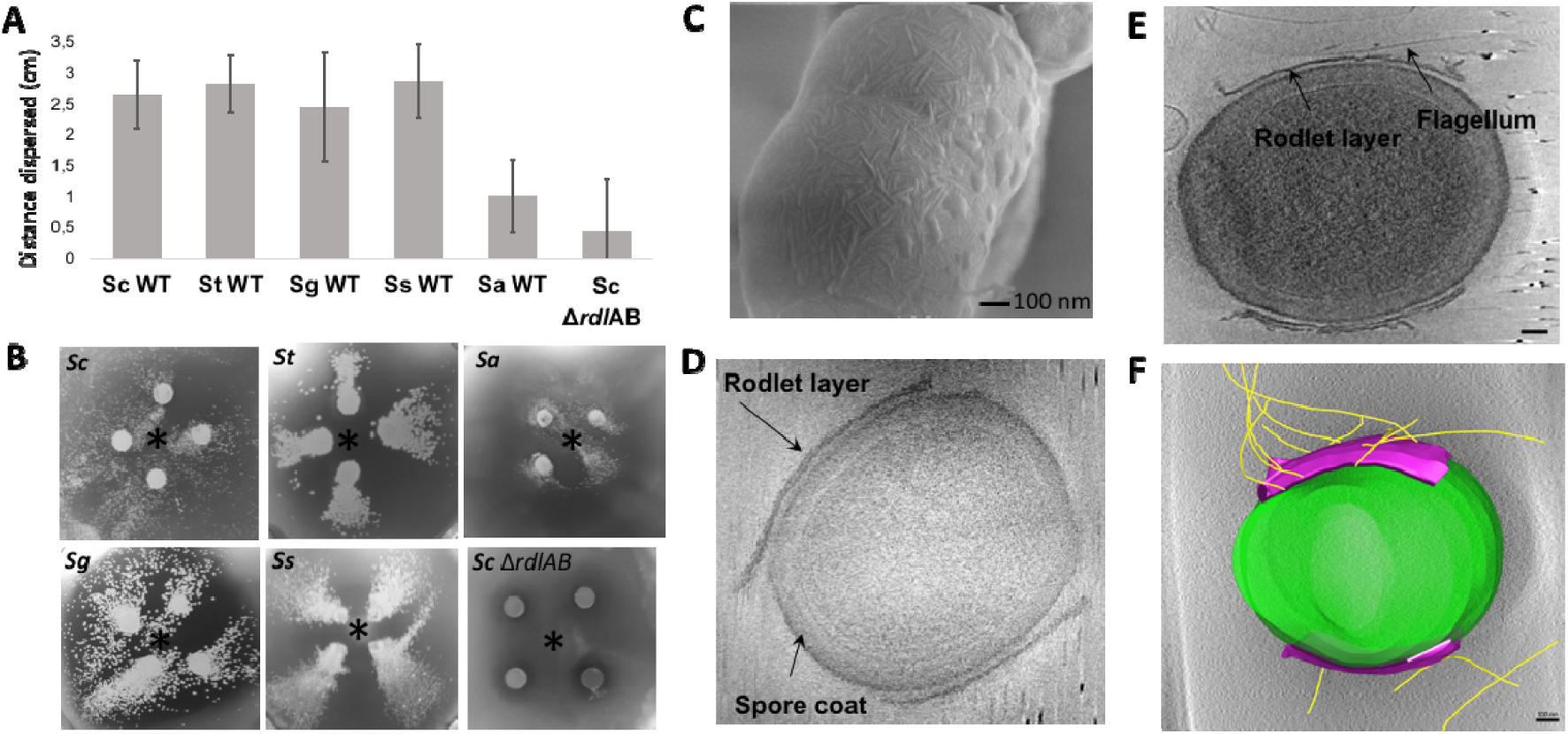
Piggy-backing of *Streptomyces* spores is facilitated by the presence of rodlin proteins. (A) *Bs* swarm assays with both wild-type (WT) *Streptomyces* spores (*S. coelicolor n* = 20, *S*. *tendae n* = 12, *S*. *griseus n* = 8, *S. scabies n* = 8, *S. avermitilis n* =12) and *Sc* spores lacking the rodlin proteins (*Sc* Δ*rdl*AB, *n* = 24) demonstrate that spores are dispersed in all tested WT species but with *S. avermitilis* dispersed the shortest distance, and dispersal is abrogated by the loss of the rodlins in *Sc*. p < 0.05 for *Sc* Δ*rdl*AB compared to WT *Sc, St, Sg*, and *Ss* strains. (B) Representative images of *Bs*/spore swarm plates from A. The *Bs* innoculation site is denoted with an asterisk. (C) SEM of WT *Sc* spores shows the rodlet layer with pair-wise rodlets of 20 nm spacing. (D) A representative cryo-ET image of isolated *Sc* WT spores shows that the rodlet layer does not cover the entire spore but leaves the poles exposed (*n* = 22). (E) Cryo-ET reconstructions show that flagella preferentially interact with the rodlet layer (*n* = 12). Scale bar 100 nm. (F) Segmentation of the reconstruction from E clearly demonstrate the flagella:rodlin interaction. Purple: rodlet layer, Yellow: flagella. Scale bar 100 nm.

### Spore dispersal by B. subtilis is facilitated by the rodlin proteins

The outer surface of most *Streptomyces* spores is characterized by a fibrillar rodlet layer, which is a striated pattern of pairwise aligned rodlets composed of the rodlin proteins^22,23^. Scanning electron microscopy (SEM) images of *Sc* and *Ss* spores show the striated rodlet layer (Fig. 3C). In previous studies the rodlets of *Sc, S. lividans, St, Sg*, and *Ss* were visually indistinguishable^23,31^. Using electron microscopy images from this and previous studies, we measured the spacing of the rodlets in these species, which is highly conserved and around ∼20 nm (when measured from the center of the rodlet fibers) (Table S1). Furthermore, the rodlin proteins from *Sc, St* and *Sg* have ∼34 % sequence identity despite the species’ distant evolutionary relation (Fig. S4)^23^.

Intriguingly, *Sa* spores are significantly less dispersed than the other *Streptomyces* species and it is the only tested species that natively lacks rodlin proteins^23^. In agreement, an *Sc* mutant strain that lacks the rodlin proteins (Δ*rdlAB*) abrogates piggy-backing by *Bs* (*n* = 3, Fig. S3). Importantly, previous studies demonstrate that the *Sc* Δ*rdlAB* strain is not delayed in germination, and does not exhibit any behavioral or phenotypic change compared to the WT strain with the exception of the rodlet layer^22,23^. In contrast, *Sc* mutants that lack proteins which produce polysaccharides found on the surface of Streptomycetes (Δ*cslA* and Δ*matAB*) are unaffected (*n* = 3, Fig. S3)^32^. However, spore dispersal was not completely abolished in the Δ*rdlAB* strain. Using identical swarm plate assays described in the section above, the Δ*rdlAB* strain is dispersed with an average maximum distance of 0.46 +/- 0.82 cm from the initial inoculation point and dispersal only occurs in 33 % of the samples (*n* = 24) (Fig. 3A,B).

To characterize how rodlins interact with flagella in three dimensions, we conducted cryo-ET experiments of samples containing *Bs* minicells and *Sc* spores. Reconstructions show that the *Sc* spores are oval shaped and possess a thick coat. The rodlet layer can be seen as a sheath around the lateral sides of the spore with frayed edges, leaving the poles exposed, and suggest that the rodlet sheath easily peels away from the cell body (*n* =22) (Fig. 3D). *Bs* flagella accumulate around and directly interact with the rodlet layer (*n* = 12) (Fig. 3E,F, <u>Movie 2</u>). However, due to the thickness of the spores the resolution is limited and we could not deduce if the flagella preferentially bind specific features of the rodlet layer. Collectively, these data suggest that the rodlet layer facilitates spore dispersal by interacting directly with flagella.

### Piggy-backing of S. coelicolor spores is not limited to Bacillus

Although *B. subtilis* is ubiquitous in soil, other genera are also flagellated and may also contribute to dispersal of *Sc* spores. We therefore conducted swarm plate assays with *Pseudomonas fluorescens* wild-type strain R1SS101, which is also attracted to plant root exudate^9^. Importantly, dispersal of *Sc* spores by *P. fluorescens* is similar to *Bs* on swarm plates (*n* = 3) (Fig. S5). These data demonstrate that piggy-backing is a widespread mechanism that allows *Streptomyces* spores to disperse at cm-scales (Fig. 4).

**Fig. 4.**
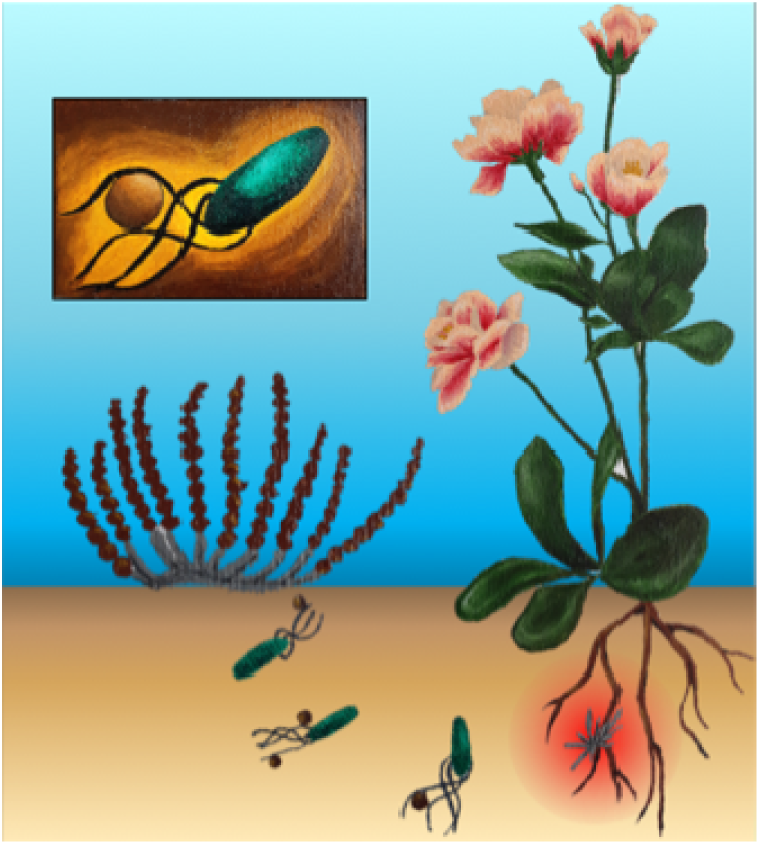
An overview of the piggy-backing model. Aerial *Streptomyces* spores are transported on the cm-scale to plant root systems by directly adhering to the flagella of motile bacteria (inset). Here, the spores germinate and produce antibiotics (red gradient) to ward off microbial competitors.

## Discussion

Sessile Streptomycetes have a complex lifecycle that involves formation of aerial hyphae that differentiate into spores. The spores of some *Streptomyces* species, including *Sc*, are dispersed over long distances by direct attachment to insects and nematodes^6^. Intriguingly, recent reports identify that specific volatile metabolites secreted by Streptomycetes attract arthropods as a mechanism for spore dispersal^7^, and can induce the formation of *Streptomyces* ‘explorer cells’ while simultaneously starving microbial competitors^33^. However, it is unclear how the spores are transported specifically at the centimeter scale to plant-root microenvironments. Here, we demonstrate that *Streptomyces* spores are able to utilize the motility machinery of motile soil microbes by directly attaching to their flagella. While these experiments demonstrate that *Sc* spores are dispersed by *Bs* and *Pf* regardless of their destination, they are both chemotactic toward favorable environments (like plant roots) and away from toxic ones. Therefore, this mechanism of dispersal, called piggy-backing, provides *Streptomyces* spores a mechanism for translocation to beneficial environments and away from harmful ones. Indeed, assays with *A. thaliana* plants demonstrate that *Bs* transports *Sc* spores to plant tissues. This allows spores to germinate near nutrient-rich plant exudate to generate filamentous colonies that produce antibiotics, thereby protecting the plant from potential phytopathogens.

Piggy-backing is facilitated by two spore coat proteins, RdlA and RdlB, which are conserved in most streptomycetes. These proteins assemble into pairwise aligned filaments, called rodlets, on the outer surface of the spores and are spaced approximately 20 nm apart. Until now, the function of the rodlets has remained elusive ^34^. Interestingly, the diameter of the bacterial flagellar filament is also ∼20 nm ^26,35^. Therefore, it’s possible that the rodlet layer provides a gripped surface for the flagella, which become ‘wrapped’ in the grooves made by the rodlin proteins and thereby facilitates spore transport. Emerging studies have demonstrated that flagella preferentially interact with hydrophobic surfaces and flagellin can undergo methylation to increase flagella hydrophobicity^36–38^. This increase in hydrophobicity allows pathogenic bacteria to adhere to host cells^36–38^, and flagellar adherence to plant cells has also been implicated in establishing colonization^39,40^. Flagella hydrophobicity may facilitate interactions with hydrophobic spores and may account for spore transport that is seen in the absence of rodlins, given that the spore surface without rodlins remains hydrophobic (in *Sa* WT and *Sc* Δ*rdl*AB strains)^22,41^. Therefore, although piggy-backing of *Streptomyces* spores is disadvantageous to motile bacteria, the same flagellar interactions that facilitate adherence to plant roots may also contribute to adherence to spores, and necessitate that motile bacteria not evolve to abrogate this interaction.

The piggy-backing model is supported by a previous study that examines the interaction of two other root-colonizing microbes: the immotile fungus *Aspergillus fumigatus* (*Af*) and the motile bacterium *Paenibacillus vortex* (*Pv*)^20^. *Af* spores are demonstrated to be dispersed by *Pv* in a swarming-dependent manner via direct attachment to flagella; dispersal is abrogated by the addition of excess purified *Pv* flagella or perturbations to the *Af* spore coat^20^. Furthermore, scanning EM micrographs show direct contact between *Pv* flagella and *Af* spores^20^. Although this study does not identify the spore coat component(s) responsible for adherence to flagella, *Aspergillus* spores also possess a rodlet layer^42^. Additionally, this study demonstrates that some *Penicillium* species are also transported by *Pv*, and these fungi possess a rodlet layer^43^. Collectively, these data may suggest that piggy-backing of spores onto motile bacteria via the formation of a striated rodlet layer is a dispersal mechanism that convergently evolved in both domains of life.

The colonization of plant roots by some Streptomycetes, including *Sc*, improves plant health and performance in a natural and sustainable manner^2–5^. Therefore, our data are applicable to industrial initiatives that aim to improve soil conditions for *Streptomyces* root colonization. Likewise, many *Aspergillus* fungi, like *Af* and *A. niger,* are human and plant pathogens. Therefore, insights into piggy-backing of these sessile organisms may elucidate unknown infection mechanisms.

---

**Table S1.** Measurements between rodlet pairs of *Streptomyces* species from electron micrographs. ^#^ In these microg raphs, the rodlin p roteins from *S. tendae* were complimented in *S. coelicolor* lacking rodlins.

| Species | Rodlet spacing | Referenced study |
| --- | --- | --- |
| *S. coelicolor* | 19.2 +/- 2.5 nm (n = 11) | This study |
| *S. griseus* | 19.2 +/- 2.1 nm (n = 12) | Claessen et al. *Mol Micro.* 2004. |
| *S. tendae*<sup>#</sup> | 21.8 +/- 4 nm (n = 10) | Claessen. et al. *Mol Micro.* 2004. |
| *S. lividans* | 20.8 +/- 1.7 nm (n = 12) | Di Berardo. et al. *J Bac.* 2008. |
| *S. scabies* | 20.9 +/- 2.2 nm (n = 10) | This study |

**Fig. S1.**
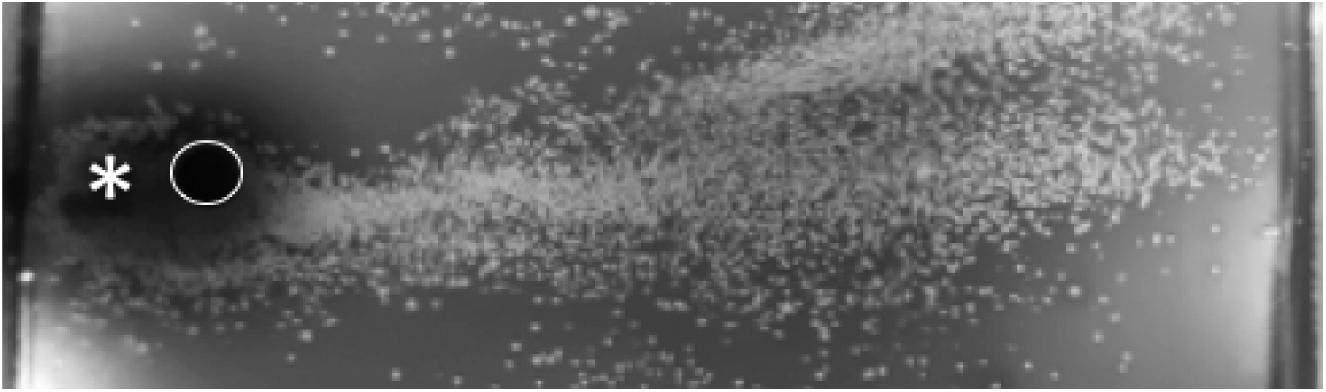
Spore dispersal assays with *Bs* and WT *Sc* on a 12 cm plate demonstrate that the spores are dispersed to the edge of the plate, which is 10 cm from the *Sc* innoculation point (*n* = 3). The *Sc* innoculation site is outlined in a white circle. The *Bs* innoculation site is denoted with an asterisk.

**Fig. S2.**
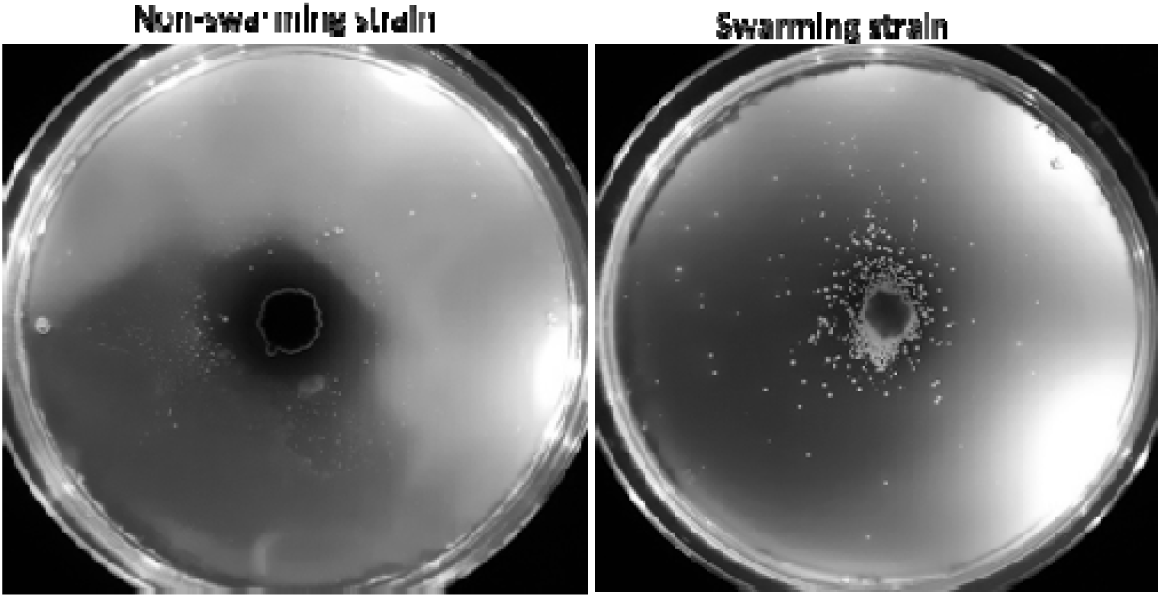
The *Bs* swarming-deficient strain does not transport *Sc* spores on 0.27 % agar plates but the swarming strain does.

**Fig. S3.**
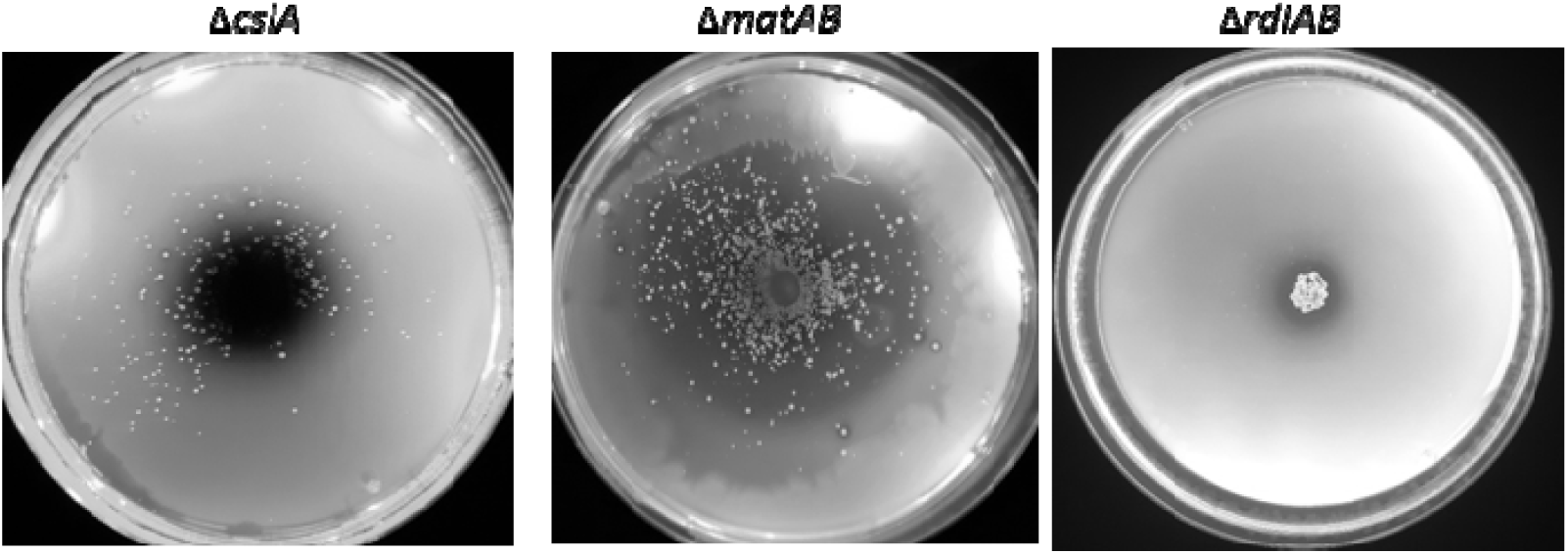
Swarm assays with *Sc* spore mutants that lack proteins which synthesize spore-coat polysaccharides (Δ*cslA* and Δ*matAB*) do not interfere with spore dispersal by *Bs* but mutants that lack the rodlin proteins (Δ*rdlAB*) interfere with dispersal.

**Fig. S4.**
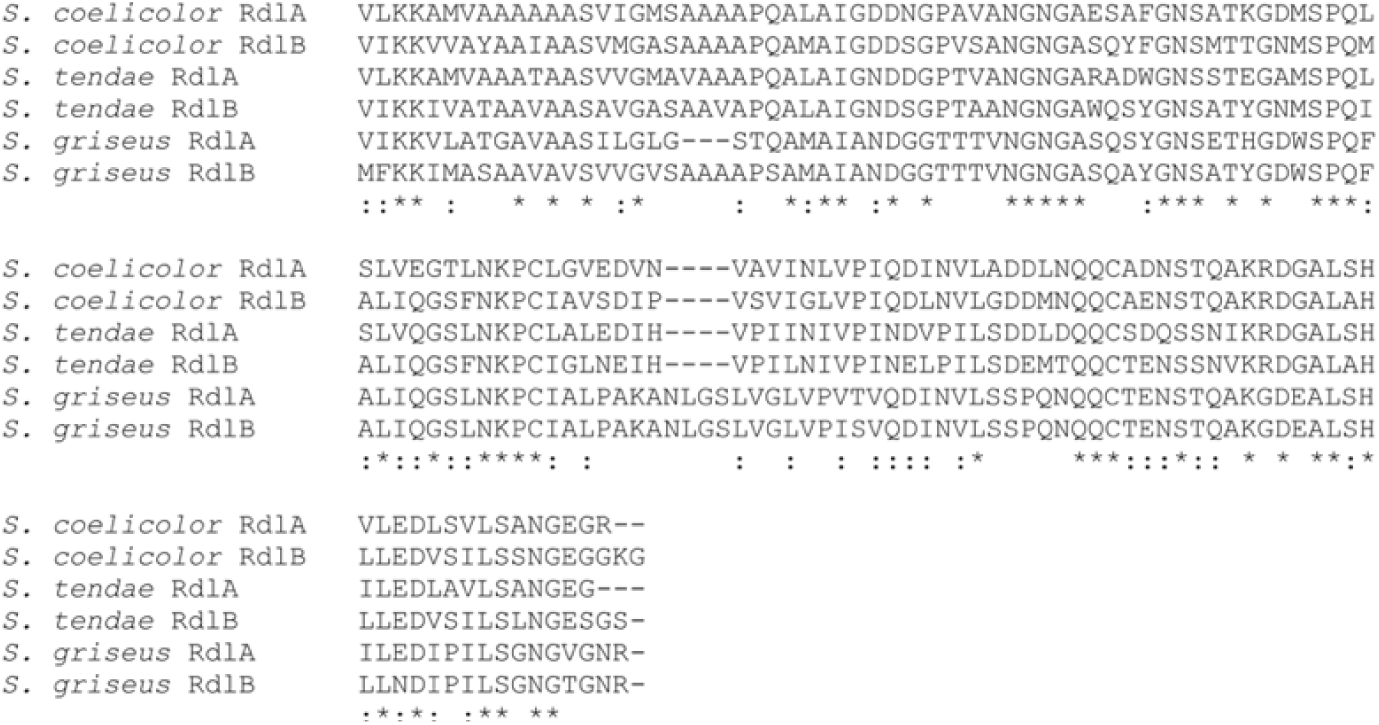
A multiple sequence alignment of the rodlins, RdlA and RdlB, from *Sc, St and Sg*. Data reproduced from Claessen *et al*. 2004.^23^

**Fig. S5.**
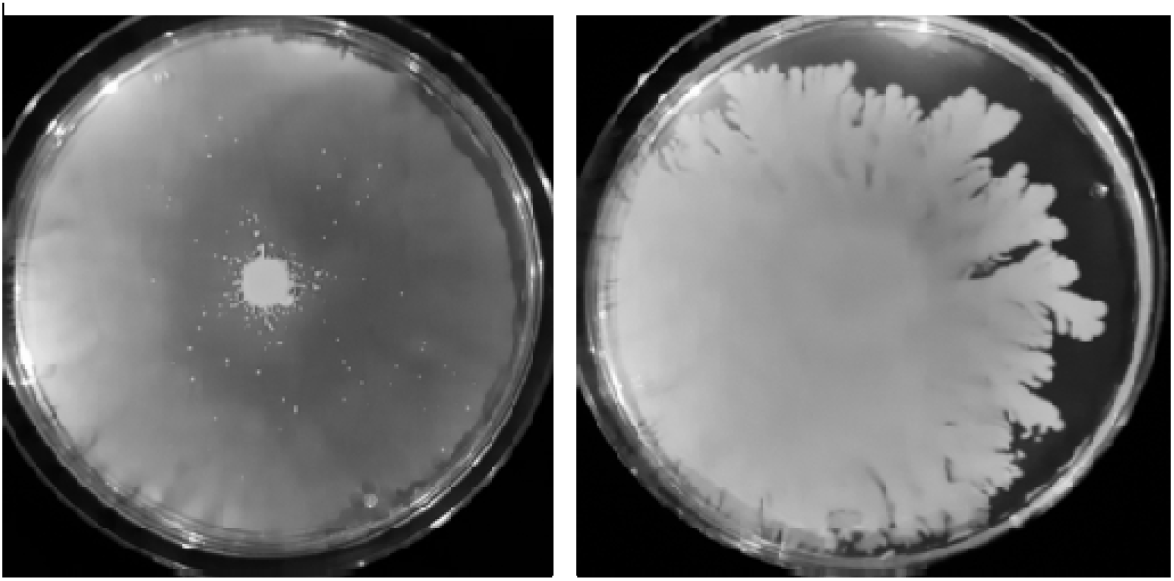
Sc spores are dispersed by *Pseudomonas fluorescens* (*Pf*). Left: *Pf* cells and *Sc* spores were both innoculated on the center of the plate. Right: *Pf* alone.

## Methods and Materials

### Streptomyces spore isolation

The following *Streptomyces* strains were used in this study: *S. coelicolor* M145^44^, *S. coelicolor* Δ*rdlAB6*^*41*^, *S. tendae* Tü901/8c^45^, *S. griseus* (ATCC 13273), *S. avermitilis* (ATCC 31267), and *S. scabies* ISP5078. Spores were harvested from MS agar plates and quantified as described before^44^.

### Bacillus subtilis cultivation

The undomesticated *Bacillus subtilis* (*Bs*) strain NCIB3610^29^ 25% glycerol stock was placed into 5 ml of LB and grown overnight at 30°C. After 16 hours of growth, 100 μL of the overnight culture was diluted into 5 ml of LB and grown at 37°C to an O.D. of 0.4-0.5.

### Pseudomonas fluorescens cultivation

The *Pseudomonas fluorescens* (*Pf*) strain R1SS101 25% glycerol stock was placed into 5 ml of 50% TB and grown overnight at 30°C. After 16 hours of growth, 100 μL of the overnight culture was diluted into 5 ml of 50% TB and grown at 37°C to an O.D. of 0.4-0.5.

### Swarm and swim plate assays

Swarm and swim plates were conducted on nutrient broth plates (0.5% peptone, 0.3% yeast extract, 0.5% NaCl) containing specific amounts of agar (0.27%-0.5%). All components were mixed, autoclaved for 20 minutes, and 30 ml of the media was poured into a plastic petri dish with a 9 cm diameter. The plates were cooled for 30 minutes in a sterile fume hood and then stored in a 4°C fridge for a maximum of 1 week. To determine if *Sc* spores are dispersed by *Bs*, 3 μL of *Bs* cells are inoculated onto the plate and 3 μL of *Sc* spore stocks are either added on the *Bs* inoculation site or to a separate inoculation site. The plates are incubated at 30°C for 5 days an imaged on a light box. The distance of spore dispersal was determined using Image J software. The correlation between *Sc* colony size and dispersal distance was quantified using Image J software with custom-made scripts.

### Fluorescence microscopy

10 μL of *Sc* spore stock was added to 1 ml of iced LB. 1 μL of the fluorescent styryl dye, FM2-10 (Thermo Fisher Scientific), was added to the 1 ml solution and inverted to mix. Excess dye was removed by rinsing the spores 4X with 1 ml of iced LB via centrifugation and decanting. After the final decantation, 1 ml of *Bs* cells with an O.D. of 0.4 were added to the spores, mixed via pipetting, and incubated at ambient temperatures for 5 minutes. Immediately before imaging, 5 μL of the samples were placed on a glass slide and a glass coverslip was placed on top. The sample was imaged on a Zeiss Axioscope A1 fluorescent microscope scope equipped with an Axiocam Mrc5 camera (Zeiss) in the Institute of Biology Microscopy Unit using a GFP filter. Images were collected and processed using Axiovision software (Zeiss).

### Cryo-electron microscopy

*B. subtilis* cells were grown to an O.D. of 0.5 and 1 ml of *B. subtilis* cells were mixed with 5 μL *S. coelicolor* spores glycerol stock and incubated at ambient temperatures for 5 minutes. Cells were concentrated by centrifugation and 3 μL aliquots of the cell suspension are applied to glow-discharged R2/2 200 mesh copper Quanti-foil grids (Quantifoil Micro Tools), the sample was pre-blotted for 30 seconds, and then blotted for 2 seconds. Grids were pre-blotted and blotted at 20 °C and at 95 % humidity. The grids were plunge-frozen in liquid ethane using an automated Leica EM GP system (Leica Microsystems) and stored in liquid nitrogen. The grids were imaged on a 120 kV Talos L120C cryo-electron microscope (Thermo Fisher Scientific) at the Netherlands Center for Electron Nanoscopy (NeCEN).

### Cryo-electron tomography

*B. subtilis* minicell strain was grown from a 20 % glycerol stock to an O.D. of 0.6 in 50 ml of LB. The cells were centrifuged at 8,000 xg for 30 minutes. The supernatant was collected and then centrifuged at 12,000 xg for 20 minutes. The resulting cell pellet was resuspended in 20 μL of LB and 8 μL of WT *Sc* spore stock was added to the cell mixture. A 1/10 dilution of protein A-treated 10-nm colloidal gold solution (Cell Microscopy Core, Utrecht University, Utrecht, The Netherlands) was added to the mixture and mixed by pipetting. The grids were prepared using an automated Leica EM GP system (Leica Microsystems) with the sample chamber set at 20 °C and at 95 % humidity. 3 μL of the sample mixture was applied to a freshly glow-discharged copper R2/2 200 grid (Quantifoil Micro Tools), pre-blotted for 30 seconds, and then blotted for 2 seconds. The grid was plunge frozen in liquid ethane and stored in liquid nitrogen.

Images were recorded with a Gatan K3 Summit direct electron detector equipped with a Gatan GIF Quantum energy filter with a slit width of 20 eV. Images were taken at a magnification of 19,500X, which corresponds to a pixel size of 4.4 Å. Tilt series were collected using SerialEM with a bidirectional dose-symmetric tilt scheme (−60° to 60°, starting from 0°) with a 2° increment. The defocus was set to – 12 μm and the cumulative exposure per tilt series was 160 e-/A^2^. Bead tracking-based tilt series alignment and drift correcting were done using IMOD^46^ and CTFplotter was used for contrast transfer function determination and correction^47^. Tomograms were reconstructed using simultaneous iterative reconstruction with iteration set to 4. Segmentation was done in IMOD.

### Plant growth

*Arabidopsis thaliana* Col-0 strain was grown from sterilized seedlings on sterilized plant MS agar media. Harvested *A. thaliana* seeds were sterilized in a sterile fume hood by incubation in 10 % bleach for 30 minutes, washed with sterile water, and then incubated in 70 % ethanol for 5 minutes. The seeds were then washed 6X with sterile water, placed on sterile filter paper, and placed in a dark 4°C fridge for 3-4 days in a sterile and parafilm-sealed petri dish. Plant agar media plates were prepared by autoclaving Murashige and Skoog (MS) media (0.22 % MS media with vitamins, 1.2 % plant agar, 0.5% sucrose, pH 5.8) and pouring 100 ml of the media into square petri dishes with 12 cm length. The plates were allowed to cool for 1 hour in a sterile fume hood. *A. thaliana* seeds were manually placed on the surface of the plates 1 cm apart by picking up the seeds with sterilized wooden picks. The plates were sealed with parafilm and placed in a climate-controlled plant growth chamber at a 20° angle so the plant roots grew on the surface of the media. The plant chamber was kept at 21°C with a 16-hour light cycle. The plants were allowed to grow for 1 month before use in chemoattraction assays (below).

### Chemotaxis attractant assays with plant roots

Chemoattraction of *Bs* cells to plant roots in the presence and absence of *Sc* spores was conducted on minimal media plates with 0.25 % agar. The media was prepared according to previous methods^48^ in round petri dishes with 9 cm diameter. 1 month old sterile *A. thaliana* plants were removed from their sterile media and placed on the edge of the minimal media plates. 3 μL of *B. subtilis* culture was placed to the center of the plate, and then 3 μL of the isolated spore stock was also added to the center. Controls of each bacteria by itself were also prepared. The plates were incubated for 16 hours at 30°C and then placed in a climate-controlled plant growth chamber for 2 weeks. The plant chamber was kept at 21°C with a 16-hour light cycle. After *Sc* colonies were visible, the plates were imaged on a light box.

## Acknowledgements

We thank Mark Landinsky at Caltech for the 3D segmentation of the *Sc* spores with *Bs* minicells, Dr. Jos Raaijmakers for the *P. fluorescens* strain R1SS101 and the *A. thaliana* Col-0 strain, Dr. Chris Rao for the undomesticated *B. subtilis* strain NCIB 3610, Dr. Daniel Kearns for the *B. subtilis* mini-cell strain minD::TnYLB kan, Dr. Rose Loria for the *S. scabies* ISP5078 strain, and Dr. Joost Willemse for assistance with the fluorescence microscopy experiments. We also thank Dr. Jos Raaijmakers and Dr. Gilles van Wezel for their on-going support and advice with this project. We thank the Netherlands Centre for Electron Nanoscopy (NeCEN) for access to cryo-EM data collection and processing facilities, and the Institute of Biology Microscopy Unit at Leiden University for access to and training with light and fluorescence microscopes. This work is part of the research programme National Roadmap for Large-Scale Research Infrastructure 2017 – 2018 with project number 184.034.014, which is financed in part by the Dutch Research Council (NWO). This project was funded by the European Union under a Marie-Sklodowska-Curie COFUND LEaDing fellowship to A.R.M.

## Notes

https://www.dropbox.com/s/gsfq798midvdcxy/Movie1.mp4?dl=0

https://www.dropbox.com/s/6j1nbdbtnp5bpqz/BS_Spore2.mrc_unweightbin.mov?dl=0

https://www.dropbox.com/s/ooy3ftlo1d62p6f/MovieS1.avi?dl=0

https://www.dropbox.com/s/8wk5ch4vrblpmgg/MovieS2.avi?dl=0

### Competing Interest Statement

The authors have declared no competing interest.

